# Mapping genetic and phenotypic diversity of *Pseudomonas aeruginosa* across clinical and environmental isolation sites

**DOI:** 10.1101/2025.09.22.677895

**Authors:** Cristina Penaranda, Evan P Brenner, Anne E Clatworthy, Lisa A Cosimi, Janani Ravi, Deborah T Hung

## Abstract

*Pseudomonas aeruginosa* is a clinically significant opportunistic pathogen adept at thriving in both host-associated and environmental settings. To define the extent to which *P. aeruginosa* isolates specialize across niches and identify genotype-phenotype correlates, we performed whole genome sequencing and comprehensive phenotypic characterization of 125 *P. aeruginosa* isolates from diverse clinical and environmental sites, evaluating virulence-associated traits, including motility, cytotoxicity, biofilm formation, pyocyanin production, and antimicrobial susceptibility. We identify that genomic diversity does not correlate with isolation source or most virulence phenotypes. Instead, we find that the two major *P. aeruginosa* clades (Groups A and B) delineate phylogeny and cytotoxicity, with Group B strains showing significantly higher cytotoxicity than Group A. Sequence analysis revealed previously uncharacterized alleles of genes encoding type III secretion effector proteins. We observed high variability amongst strains and isolation sources in all four assayed virulence phenotypes. Antimicrobial resistance (AMR) is exclusively observed in clinical isolates, not environmental, reflecting antibiotic exposure-driven selection. Bacterial GWAS revealed a statistically significant association between cytotoxicity and *exoU* presence, and we identified a novel *exoU* allelic variant with decreased cytotoxicity, demonstrating that functional diversity within well-characterized virulence factors may still influence pathogenic outcomes. In summary, our analyses of 125 diverse isolates suggest that the ability of *P. aeruginosa* to thrive across diverse niches is driven by broadly conserved genetic repertoire rather than niche-specific accessory genes.

**Importance:** *Pseudomonas aeruginosa* is a clinically significant opportunistic pathogen adept at thriving in both host-associated and environmental niches. A major gap in our understanding of this difficult-to-treat pathogen is whether niche specialization occurs in the context of human disease. Addressing this question is critical for guiding effective infection control strategies. Previous large-scale studies have focused solely on genotypic or phenotypic analyses; when paired, they have been limited to a single phenotypic assay or to a small number of isolates from one source, or relied on PCR-based methods targeting a restricted set of genes. To comprehensively uncover niche specialization and pathogenic versatility, we performed whole genome sequencing and phenotypic characterization of 5 virulence-associated traits including AST of 125 clinical and environmental *P. aeruginosa* isolates. Our systems-level findings challenge reductionist models of bacterial niche specialization, instead supporting an integrated view where conserved genomic systems enable opportunistic pathogenesis across diverse environments.

## Introduction

*Pseudomonas aeruginosa* is a ubiquitous environmental Gram-negative bacterium capable of both acute and chronic infections at mucosal surfaces such as urinary and respiratory tracts, skin, and eyes. *P. aeruginosa* is a leading cause of nosocomial infections, especially in people with cystic fibrosis (CF). Importantly, high intrinsic antimicrobial resistance (AMR) complicates *P. aeruginosa* treatment^1^. Multidrug-resistant (MDR) *P. aeruginosa* in the US caused an estimated 2,700 deaths in 2017 and was designated a CDC threat level of “serious” in 2019^2^.

To understand the pathogenic versatility of *P. aeruginosa*, it is essential to consider the genetic repertoire across the species. First sequenced 25 years ago^3^, over 40,000 *P. aeruginosa* genomes have been deposited to NCBI as of 2025. Compared to other bacterial species, the pangenome of *P. aeruginosa*, composed of conserved core genes and variable accessory genes, is large and has high genetic complexity^4^. The conservation of core metabolic and virulence genes across environmental and clinical isolates is thought to explain the ability of this bacterium to survive in diverse ecological niches and host tissues^5,6^. Nevertheless, niche specialization in the context of human disease has not been defined and will be critical to effectively focus infection control efforts.

Combining comparative genomic analysis with phenotypic characterization allows the correlation of specific genotypes to phenotypic differences. It provides insights into the complex biology underlying pathogen evolution and diversification through the identification of conserved and novel genetic elements within a species^7^. Advances in DNA sequencing technologies now permit fast and inexpensive sequencing of bacterial genomes, opening the door for methods standardly used in human population genetics, like genome-wide association studies (GWAS), to better understand bacterial genetic flow and evolution during infection^8,9^. For instance, analysis of *Campylobacter* isolates revealed that genes encoding vitamin B5 biosynthesis are associated with host specificity^10^. Analysis of within-species bacterial strain variation provides insights into the exchange of genes through horizontal gene transfer (HGT) and the evolution of genes through changes at the nucleotide level^7^. The impacts of such changes on pathogenesis are of great interest as they may reveal the mechanisms of host-pathogen interactions and uncover novel therapeutic targets. While genomic analysis has yielded significant insights into *P. aeruginosa* population structure and diversity, bridging the gap between sequenced genotypes and resulting phenotypes, remains challenging.

We report the whole genome sequencing and comprehensive phenotypic characterization of 125 *P. aeruginosa* strains, including environmental isolates and clinical isolates that represent the major body sites that this pathogen colonizes (lung, skin, urinary tract, blood, eye). Phenotypic characterization in four virulence-associated assays, including cytotoxicity, motility, biofilm, and pyocyanin production, as well as antimicrobial resistance (AMR) status for clinically relevant antibiotics (gentamicin, tobramycin, amikacin, imipenem, ceftazidime, piperacillin, levofloxacin, and ciprofloxacin) (**Figure 1**). Our results recapitulate the previously identified phylogenetic separation between Groups A and B; our novel analysis of isolate source identified no discernible associations between phylogeny and isolation source. Moreover, there were no distinct associations between isolation source and virulence in the phenotypes assayed. We observed that AMR was frequent in clinical strains, and nearly absent in environmental isolates, supporting antibiotic exposure-driven clinical AMR. Sequence analysis of the genes encoding type III effector proteins revealed previously unrecognized variability, including alleles that are specific to each phylogenetic Group. Our findings highlight the sheer diversity within the *P. aeruginosa* population, genetically and phenotypically.

**Figure 1.**
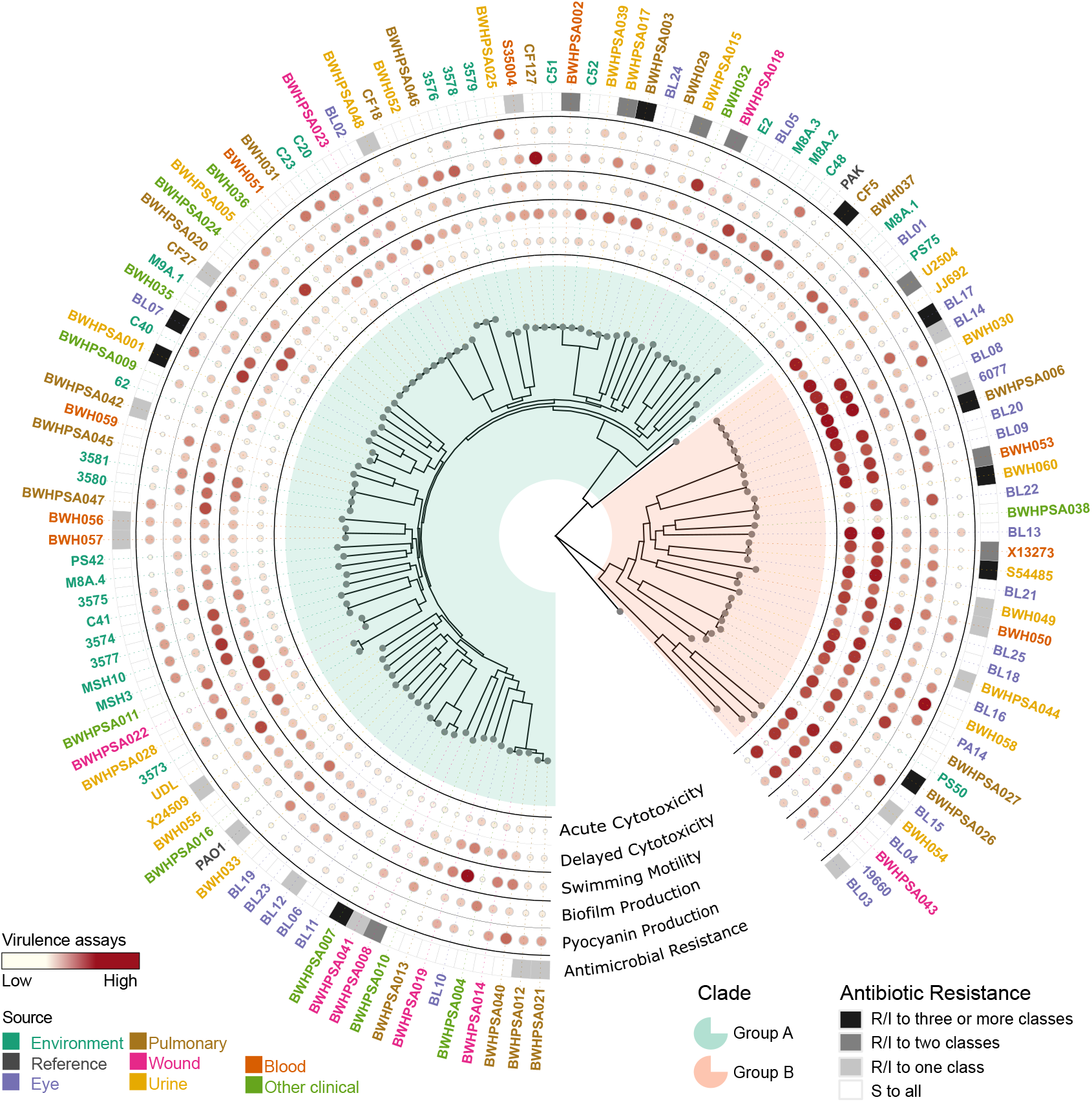
Summary of phylogenetic and phenotypic data of newly sequenced *Pseudomonas aeruginosa* strains. Whole genome sequencing and phenotypic characterization of 125 clinical and environmental strains and 3 lab reference strains (PAO1, PAK, and PA14).

Collectively, these data suggest that the large genetic repertoire of *P. aeruginosa* grants the versatility to cause infection across body sites, regardless of origin.

## Materials and Methods

### Strains

BWH/BWHPSA strains were collected by the Clinical Microbiology Laboratory at Brigham and Women’s Hospital (Boston, MA). Collection of discarded, de-identified bacterial strains was approved as non-human subjects research by the Partners Healthcare Institutional Review Board (2009P001837). Eye (BL) strains were obtained from Wolfgang Haas (Bausch & Lomb). Environmental strains were obtained from Roberto Kolter, Harvard Medical School (Boston, MA), and Paula Suarez, Simon Bolivar University (Venezuela). Other strains were obtained from Frederick M. Ausubel, Massachusetts General Hospital (Boston, MA), and Steve Lory, Harvard University (Boston, MA). We aimed to represent a diversity of infections since previous analysis has focused mainly on strains isolated from the lungs of PwCF. Our collection of clinical strains includes 17 strains previously analyzed by Wolfgang *et al*. by microarray^6^, as well as 59 new strains collected at Brigham and Women’s Hospital (Boston, MA) and 25 eye isolates provided by Bausch & Lomb. The latter set was chosen to provide geographical diversity to our collection. Importantly, the strains collected at Brigham and Women’s Hospital were only passaged once before sequencing and phenotyping with minimal opportunity for mutations to be acquired outside of the host. For our analysis, we also included the previously genome-sequenced laboratory strains PAO1^3^, PA14^11^, and PAK^12^. Metadata, including isolation source, are presented in **Supplemental Data 1**.

### Whole genome sequencing

Strains were grown overnight in lysogeny broth (LB). Genomic DNA was purified using the DNeasy Blood & Tissue kit (Qiagen) per the manufacturer’s protocol. Whole genome sequencing was performed at the Broad Institute Genomic Core using paired-end sequencing. Libraries were sequenced on the Illumina HiSeq 2000 platform and sequencing runs deposited in the Sequence Read Archive (SRA) (**Supplemental Data 1**).

### Genome assembly and analysis

For assembly, paired-end reads for all 125 successfully sequenced isolates were downloaded from NCBI SRA using SRA Toolkit^13^ (v3.0.0) to the University of Colorado Alpine HPC cluster. Reads were processed by FASTP^14^ (v1.0.1) for QC and adapter trimming, then assembled *de novo* with Shovill^15^ (v1.1.0). Genome annotation was completed for all samples by Bakta^16^ (v1.9.4) using database version 5.1 and a Prodigal training file using *P. aeruginosa* PAO1. Genome assembly and annotation completeness were assessed through QUAST^17^ (v5.3.0) and BUSCO^18^ (v5.8.3) with BUSCO dataset odb12^19^, respectively.

To explore variation across our isolates plus three reference genomes, PAO1, PA14, and PAK, Panaroo^20^ (v1.5.2) was used to generate a 128-strain pangenome. After processing, pseudogenes were removed for downstream analysis. Scoary2 (v0.0.15) was used for measuring statistical associations between gene presence and phenotypic traits, using the pangenome gene presence/absence matrix, and a phylogenetic tree that was produced by IQ-TREE2 as described below. Finer variation was extracted using Snippy^21^ (v4.6.0) and the PAO1 reference to extract polymorphisms across all isolates. For analysis within phylogenetically distinct Group B isolates, Snippy was run a second time against the Group B PA14 reference to explore polymorphisms in Group B-specific genes like *exoU*.

### Phylogenetic analysis

All called single-nucleotide polymorphisms against PAO1 were used as the core SNP set to infer a phylogenetic tree with IQ-TREE2^22^ (v2.4.0), with the best-performing substitution rate model, GTR+F+I+R6, chosen through ModelFinder^23^ (v0.1.7) (**Supplemental Data 2**). The resulting phylogenetic tree was tested using 1000 SH-like aLRT replicates^24,25^ implemented in IQ-TREE (**Supplemental Data 2, 3**). The phylogenetic tree was visualized and figures generated through R (v4.4.1) in RStudio (v2025.05.1), and R packages BiocManager^26^ (v3.19), ggplot2^27^ (v3.5.2), ggtree^28^ (v3.12.0), ggtreeExtra^29^ (v1.14.0), ggnewscale^30^ (v0.5.1), ape^31^ (v5.8.1), tidyr^32^ (v1.3.1), dplyr^33^ (v1.1.4), RColorBrewer (v1.1.3), ggsignif^34^ (v.0.6.4), and svglite^35^ (v2.2.1).

### Resistance gene analysis

Known resistance-associated genes were identified in assemblies using AMRFinderPlus^36^ (v4.0.3) with database 2025-06-03.1.

### Swimming assay

Swimming motility was measured on 0.35% agar plates as previously described with minor modifications^37^. Briefly, a sterile toothpick was used to pick individual colonies grown on LB plates and stabbed into the agar layer of a fresh 0.35% agar plate. Plates were incubated upright for 16– 17hrs at 30°C. Images were acquired on a gel imaging station. The diameter of the swimming zone was measured using ImageJ^38^. Each strain was assayed at least twice, and the average swimming zone was calculated. Phenotypic results are available in **Supplemental Data 1** for all assays.

### Biofilm assay

Biofilm formation was measured using crystal violet as previously described^39^. Log-phase bacteria grown in LB (OD_600_= ∼0.3) were diluted to 2.5×10^6 CFU/mL and plated in tissue culture-treated 96-well plates at 100μL/well as 8 replicates. Plates were incubated for 24hrs at 30°C. Plates were washed in water and stained with 200μL/well 0.1% crystal violet, incubated for 30min at RT, and washed three more times in water. 100μL/well acetic acid (33%) was added, and absorbance was measured at 590nm. Strains were assayed two or three times. All technical and biological replicates were averaged.

### Pyocyanin assay

Pyocyanin was extracted from the supernatant fraction of strains grown overnight in LB as previously described^40^. All strains were assayed in triplicate.

### Epithelial cell cytotoxicity assay

Host cell viability was measured using CellTiterGlo at 4hrs (acute cytotoxicity) and 24hrs (delayed cytotoxicity) post-infection in a gentamicin protection assay. Briefly, 5637 human bladder epithelial cells (RRID: CVCL_0126) were seeded overnight in RPMI media supplemented with 10% FBS. Log-phase bacteria grown in LB (OD_600_ = ∼0.3) and diluted in media were used to infect cells at MOI=1 in triplicate and centrifuged to synchronize infection. Cells were incubated for 2hrs, extracellular bacteria were removed, media containing 200μg/mL gentamicin was added, and incubated for an additional 2hrs and cell viability was measured at 4hrs. For measurement of delayed cytotoxicity, the media were replaced with media containing 25μg/mL gentamicin to prevent reinfection, and cell viability was measured at 24hrs. The assay was repeated at least three times for most of the strains. Some strains could not be assayed for delayed cytotoxicity due to gentamicin resistance (shown as “n/a”).

### Antibiotic susceptibility assay

Antibiotic susceptibility was assayed using BBL Sensi-Discs (BD Diagnostics): ciprofloxacin (5µg), ceftazidime (30µg), imipenem (10µg), piperacillin (100µg), levofloxacin (5µg), tobramycin (10µg), gentamicin (10µg), and amikacin (30µg). Muller-Hinton agar plates were inoculated with overnight bacterial cultures grown in LB. Plates had 4-5 discs placed and were incubated 16hrs at 37°C. Images were acquired on a gel imaging station, and the diameter of the zone of inhibition was measured using ImageJ. Most strains were assayed at least twice per antibiotic, and the average zone of inhibition was calculated. Susceptibility was determined based on the manufacturer’s Zone Diameter Interpretive Chart for *P. aeruginosa* reference strain ATCC 27853.

### *exoU* allele cloning and overexpression

Alleles of *exoU* were amplified from PA14 and BL18 (using 5’ TTCGGTACC ATGCATATCCAATCGTTGG and 5’ ATTAAGCTTTCAAACGAACACTAACGC primers) and cloned into the pHERD20T vector using KpnI and HindIII. Plasmids were transformed into a PA14Δ*exoU* strain. Epithelial cell cytotoxicity of the overexpressing strains was assayed as previously described^41^.

## Results

### Phylogenetic diversity

Advances in sequencing technologies permit fast and inexpensive sequencing of bacterial genomes to better map bacterial genetics to evolution in the environment and during infection^42,43^. To gain insights into the pathogenesis of *Pseudomonas aeruginosa*, we assembled a diverse collection of clinical isolates (97) from blood (9), eye (27), pulmonary (20 for people with (4) and without (16) CF), urine (22), wound (8) and other infections including ear infections and abscesses (11), as well as environmental isolates (28) (**Supplemental Figure 1**). After paired-end Illumina whole genome sequencing, we performed *de novo* genome assembly on all 125 strains. The number of contigs for the final assemblies ranged from 76 to 425. The average genome size was 6.65 Mbp, with an average CDS count of 6074 (**Supplemental Figure 1**). We constructed phylogenetic trees that included the reference strains PAO1, PA14, and PAK based on the entire concatenated SNP set extracted across all genomes. This set of 128 strains, including the 125 newly sequenced strains and the 3 reference strains, was used for all subsequent analyses.

Recapitulating prior analyses, the population structure consists of two major clades^7,44^: 92 strains, including the reference strains PAO1 and PAK, belong to Group A, while 34 strains, including PA14, belong to Group B (**Figure 2A, Supplemental Figure 2A**). Two strains (PS75 and BL03) group separately from the major clades, potentially belonging to minor clades as has been previously suggested^44^. None were phylogenetically related to the taxonomic outlier PA7 (**Supplemental Figure 2A**)^8^. Analysis by Ozer *et al* of intra-versus inter-group recombination events suggested that Groups A and B inhabit distinct ecological niches^44^; our results only support this conclusion for environmental strains, which are overrepresented among Group A isolates (**Supplemental Figure 2B, 2C**). However, all other clinical strains were evenly divided between Groups A and B.

**Figure 2.**
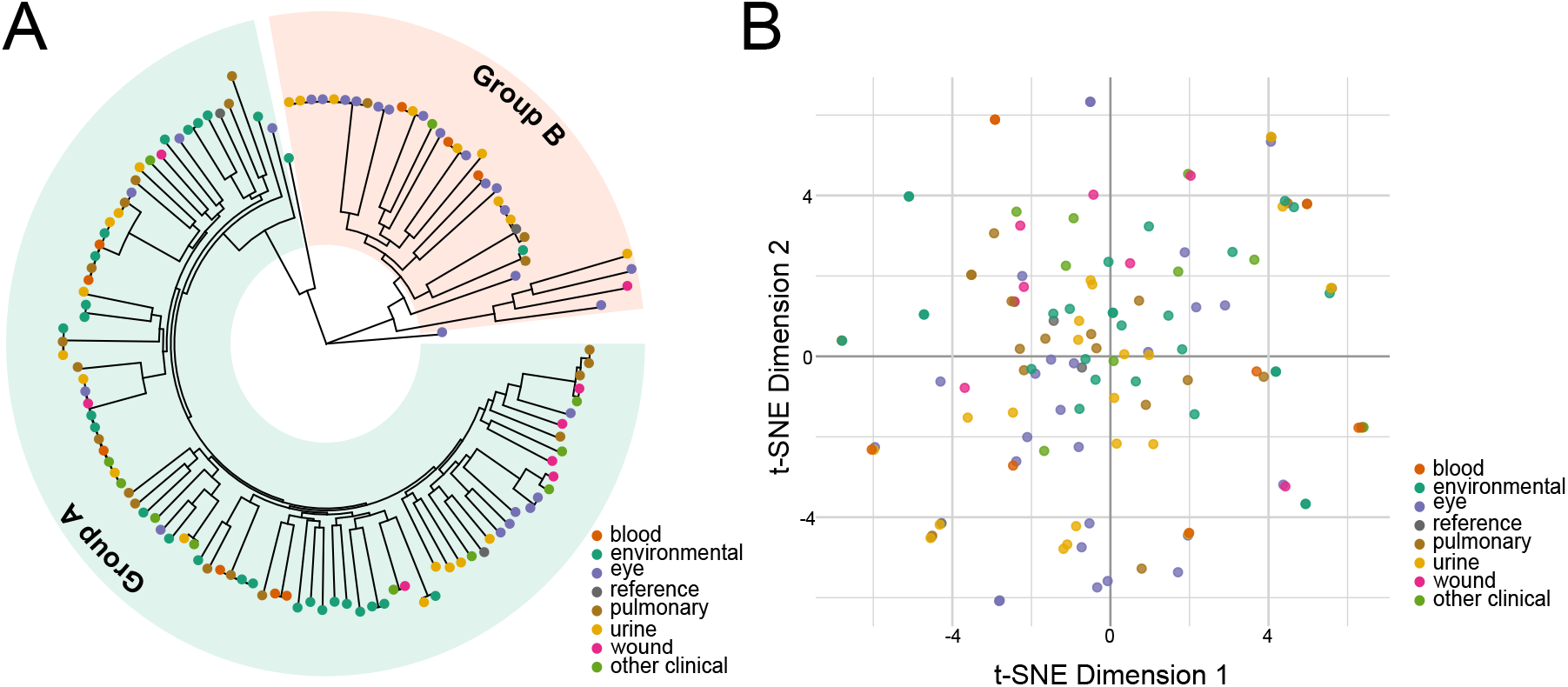
Phylogenetic distribution of newly sequenced *Pseudomonas aeruginosa* strains. **A**. Circular phylogenetic tree based on nucleotide positions with sequencing coverage across all strains. Strains are colored at the tree tips by isolation source, and broader phylogenetic groups are highlighted. **B**. t-SNE representation of the accessory genome. Genes present in <95% of strains were considered accessory and used for this analysis. The isolation source had little correlation with phylogeny and accessory genome content.

Strains did not cluster phylogenetically based on their anatomic site of infection (**Figure 2A**), in line with previous population-wide analysis^6^. We also analyzed the accessory genome by examining the gene presence/absence patterns per genome by t-SNE to determine whether strain-specific accessory genes correlated with anatomic site of infection; we did not see evidence of isolation source correlating with pangenomic accessory gene content (**Figure 2B**). No statistically associated genes were found between isolation sources (**Supplemental Data 4**).

### Sequence analysis of type III effectors

*P. aeruginosa* encodes four type III secretion system (T3SS) effector proteins (ExoU, ExoT, ExoS, and ExoY) that target the eukaryotic membrane and cytoskeleton and cause cytotoxicity^45^. The prevalence of the four effector genes *exoY, exoT, exoS*, and *exoU* in clinical and environmental strains has been widely studied due to their critical role in disease^7,46,47^. In agreement with previous reports^46^, we found that the *exoS* or *exoU* genes are mutually exclusive: 93 strains carry intact *exoS*, while the remaining 33 strains carry *exoU* (**Figure 3**). Furthermore, all 128 strains carry the *exoT* gene, supporting that this is not a variable trait^47^. We identified a frameshift mutation in CF5 (*exoT* I182fs) that likely results in a non-functional protein (**Supplemental Data 5**). Therefore, although all strains carry the *exoT* gene, not all strains appear to have a functional ExoT protein.

**Figure 3.**
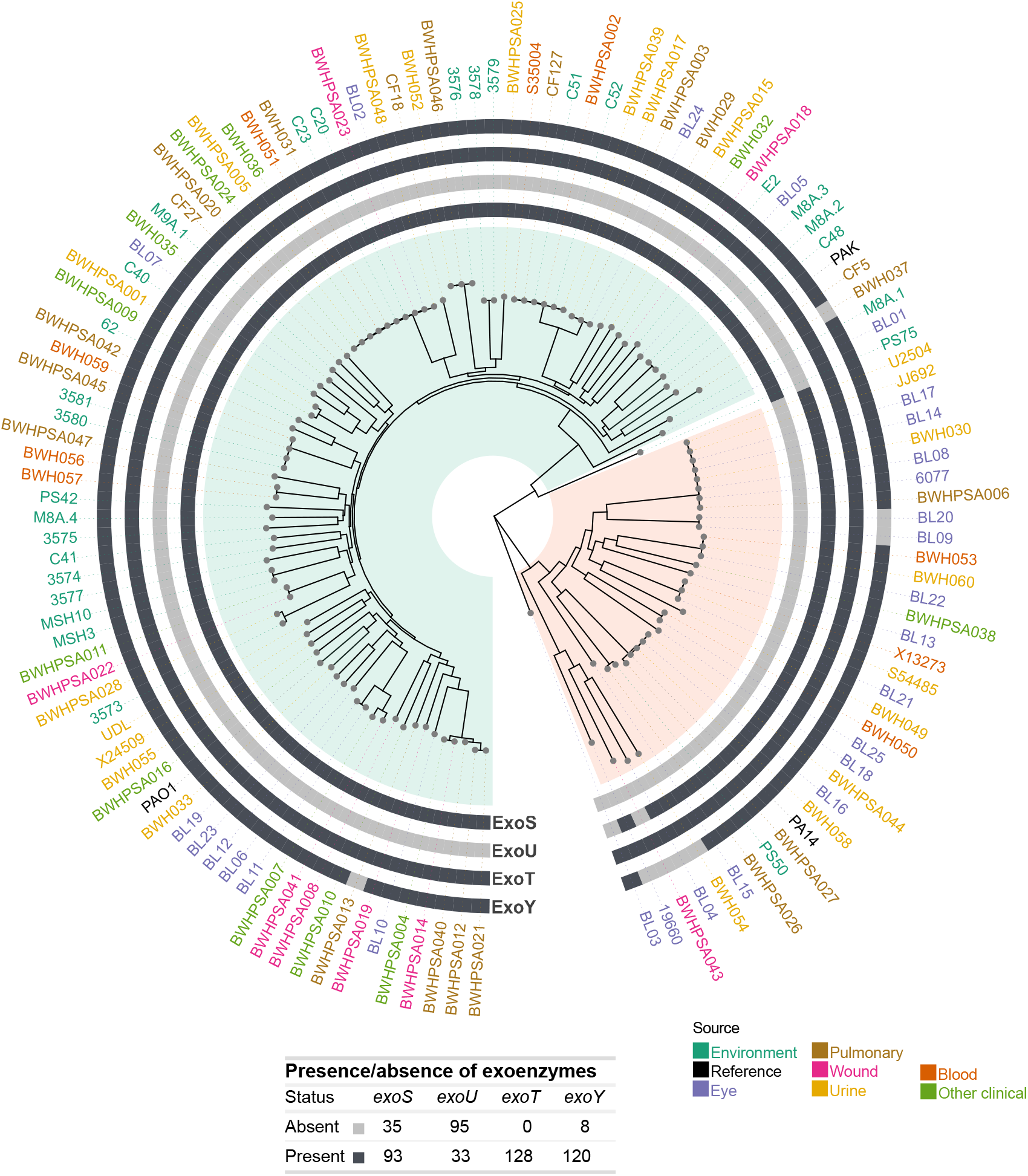
Genomic analysis of type III effectors. Circular phylogenetic tree showing the presence or absence of the four type III effectors (*exoY, exoU, exoT, exoY*) in the outer rings for each strain. Major Groups A and B are highlighted in green and orange, respectively. The presence or absence of type III effectors across strains is tabulated. Strain names are colored by isolation source, as in **Figure 1**.

Comparative analysis of all effector gene sequences revealed wide variation within the population (**Supplemental Data 5**). As has been previously described, the PA14 version of ExoY has an altered C terminus compared to PAO1, resulting from a frameshift mutation (F374fs) that adds 36 amino acids^48,49^. Of the 120 strains that carry an *exoY* gene, 7 isolates encoded either a 410aa (n = 3) or 414aa (n = 5, including PA14) protein, while 92 (including PAO1) isolates encoded the PAO1-like 378aa protein. All strains carrying a PA14-like *exoY* sequence belong to Group B, suggesting that this frameshift mutation arose after the two phylogenetic groups diverged. We also identified 5 additional frameshift or premature stop mutations that are predicted to result in proteins of different lengths, which have also been previously reported^50^. Strains C51 (Tyr186*), 3575 (Asn12fs), and BWHPSA015 (Trp359fs) each showed Group A frameshifts resulting in protein lengths unique to that strain. Finally, Leu243Fs was seen in 10 Group B isolates, yielding a predicted 248aa variant not annotated as ExoY by Bakta, but clustered with ExoY sequences by CD-HIT in Panaroo.

Additionally, we observed a novel polymorphism in the C terminus of the *exoT* gene that is predicted to encode a protein of different length (**Supplemental Data 5**). The PAO1 sequence encodes a 457aa-long protein and is shared by 120 strains, while only 4 strains share the PA14 SNP (Gln443*), resulting in a shorter 442aa-long protein. There was a frameshift mutation in one strain (CF5 G181fs) that is also predicted to result in a shorter protein in Group A.

For *exoU* or *exoY*, we did not observe alleles predicted to result in proteins of different lengths.

### Phenotypic diversity

To understand whether virulence phenotypes or AMR correlate with isolation source and phylogeny, we characterized all strains with four functional *in vitro* virulence assays: 1) production of pyocyanin, a phenazine compound that induces eukaryotic cell oxidative stress^9^, 2) flagellum-dependent swimming motility^37^, 3) formation of biofilms, associated with nosocomial infections^51^, and 4) acute (4hr) and delayed (24hr) host cell cytotoxicity, primarily attributable to injection of effector proteins into the host cytosol by the type III secretion system (T3SS)^52^. Additionally, we characterized AMR to eight antibiotics spanning five major classes: quinolones (ciprofloxacin, levofloxacin), aminoglycosides (tobramycin, gentamicin, amikacin), cephalosporins (ceftazidime), carbapenems (imipenem), and ß-lactamase inhibitors (piperacillin).

There was high variability amongst strains and isolation sources in the four virulence phenotypes assayed (**Figure 4A–E**). Of note, environmental strains were significantly more motile than pulmonary or clinical strains from other sources, which showed the least motility. Eye strains demonstrated significantly lower host cell survival (increased cytotoxicity) and increased biofilm production than pulmonary and environmental strains. Pulmonary strains included both CF (n=4) and non-CF (n=16) isolates; however, these low sample numbers did not provide sufficient statistical power for genotype-phenotype analyses.

**Figure 4.**
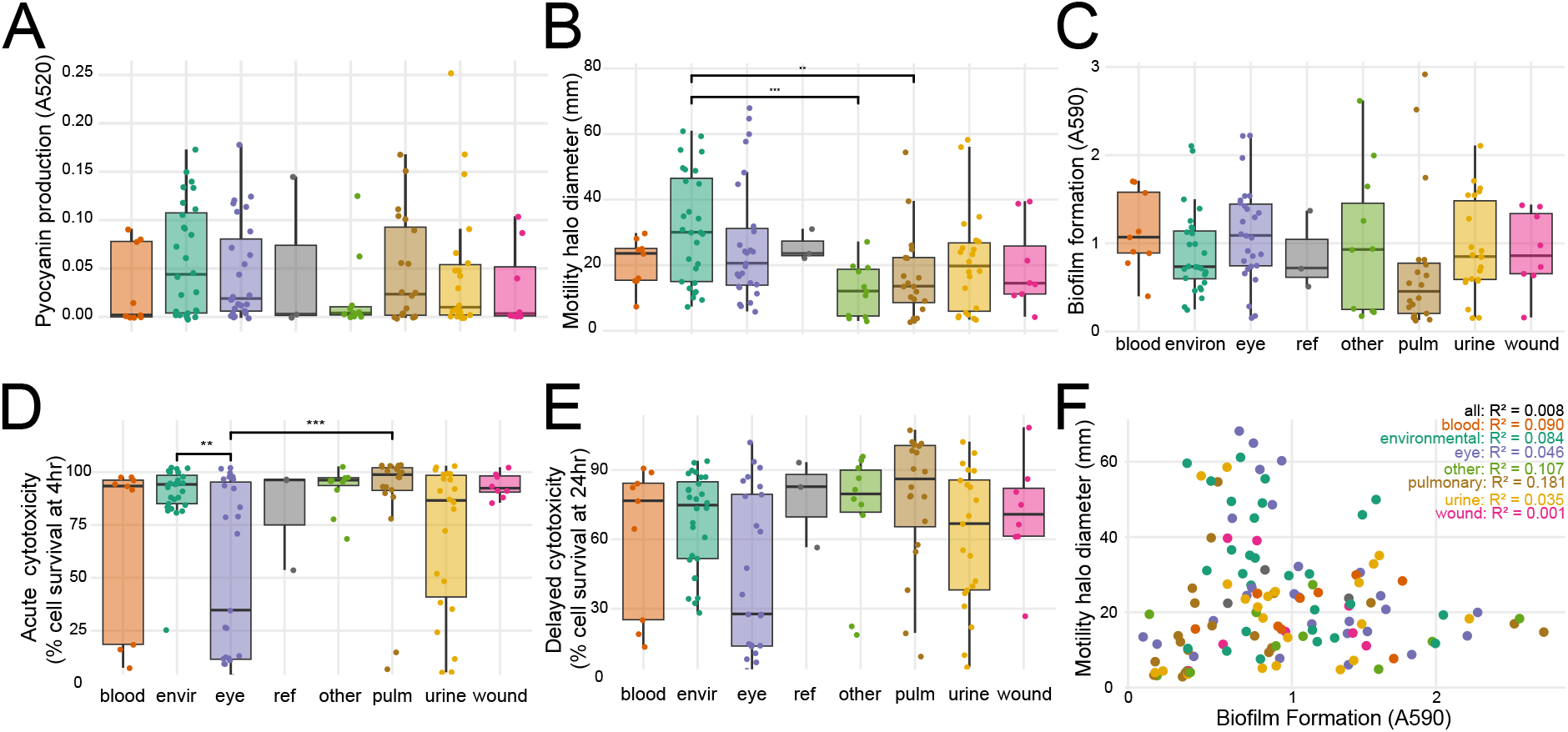
Phenotypic diversity of newly sequenced *Pseudomonas aeruginosa* strains. **A**. Pyocyanin production in the supernatant fraction of overnight LB cultures. **B**. Swimming motility on 0.35% agar plates. **C**. Biofilm formation measured by crystal violet staining. **D**. Acute cytotoxicity of human bladder epithelial cells at 4 hrs post-infection. **E**. Delayed cytotoxicity of human bladder epithelial cells at 24 hrs post-infection. **F**. Correlation of swimming motility and biofilm formation per strain. Spearman R^2^ for each isolation group is shown. For A–E, Wilcoxon test with Benjamini-Hochberg correction for multiple testing: ** p<0.01; *** p<0.001.

Biofilm formation and motility have been previously reported to be positively correlated, and it has been hypothesized that movement towards and attachment to a surface are required for biofilm formation^53,54^. However, we did not see a correlation between biofilm formation and swimming motility either for the entire population or when analyzed for each isolation site individually (**Figure 3F, Supplemental Data 6**). It is nevertheless possible that other types of motility, such as twitching or swarming motility, are required for biofilm formation.

Within our strain collection, antibiotic resistance was widespread for clinical isolates but absent for environmental isolates (**Figure 5A, Supplemental Figure 4**). Resistance to at least one antibiotic was observed in 36 of 97 clinical strains, 18 strains were resistant to at least 2 drug classes, and 10 showed clinical multidrug resistance (defined by resistance/intermediate resistance to 3 or more antibiotic classes), highlighting the overall high level of AMR seen in the population (**Figure 5B**). Resistant strains did not cluster phylogenetically (**Supplemental Figure 4)**. None of the 28 environmental strains showed any resistance. This aligns with prior observations^55^ and supports the idea that the high incidence of antibiotic resistance among clinical strains is likely due to selection pressure from treatment of *P. aeruginosa* infections.

**Figure 5.**
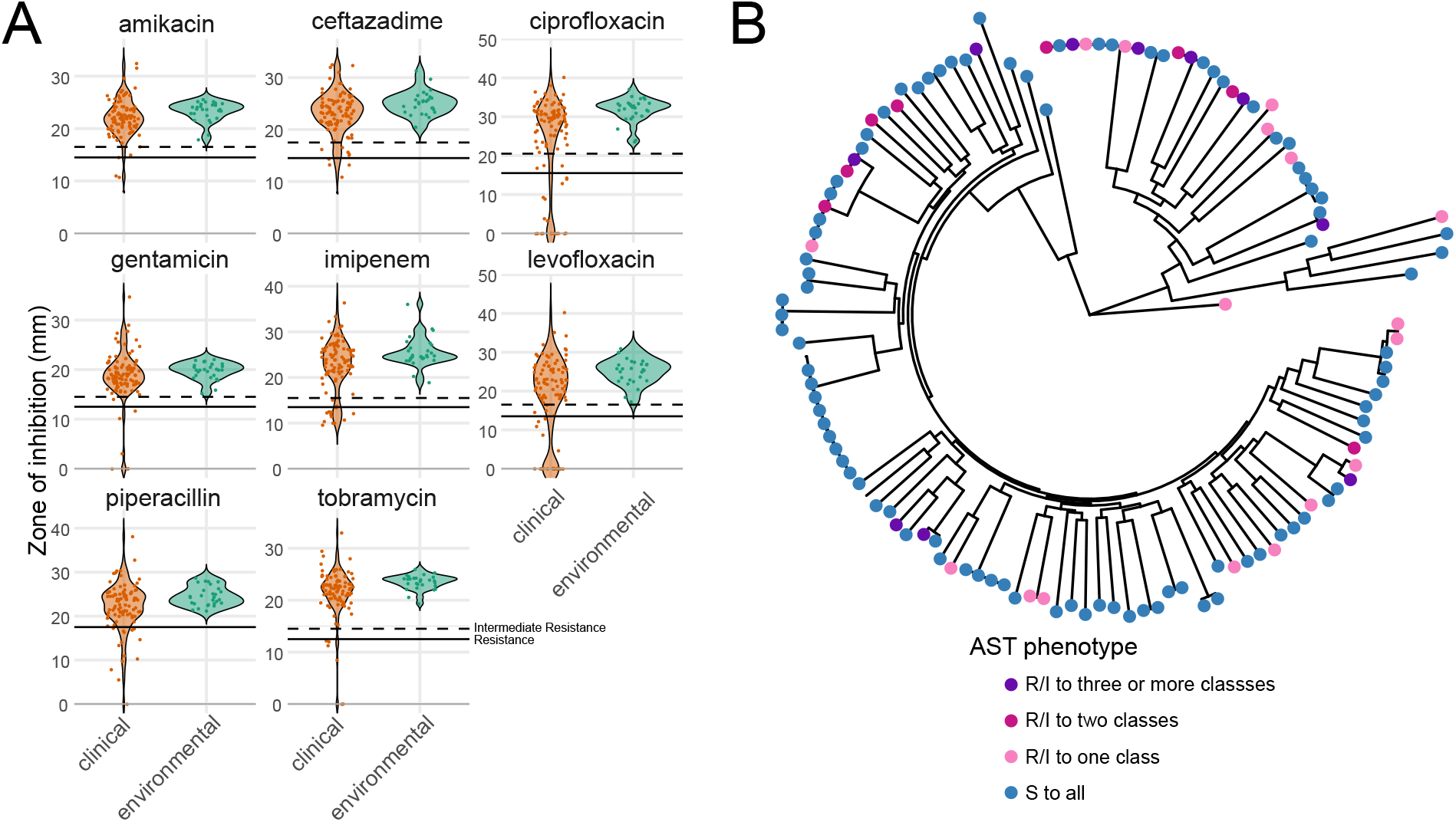
Antibiotic susceptibility of *Pseudomonas aeruginosa* strains. **A**. Zone of inhibition for each strain to eight antibiotics representing five classes commonly used to treat *P. aeruginosa* infections: quinolones (ciprofloxacin, levofloxacin), aminoglycosides (tobramycin, gentamicin, amikacin), cephalosporins (ceftazidime), carbapenems (imipenem), and β-lactamase inhibitors (piperacillin). Solid lines indicate resistance and dashed lines indicate intermediate resistance as based on the Zone Diameter Interpretive Chart for *P. aeruginosa* reference strain ATCC 27853. Clinical strains shown in orange; environmental strains shown in green. **B**. Circular phylogenetic tree as in **Figure 1A** showing AST phenotype.

We were able to identify specific genetic determinants underlying antibiotic resistance phenotypes for most resistant strains using AMRFinderPlus (**Supplemental Data 7**). Amikacin and gentamicin resistance in BWHPSA008, and levofloxacin resistance in BWHPSA044 and BWH050 did not have distinct resistance-conferring mechanisms. For these exceptions, AMRFinderPlus did identify genes generally linked to resistance, but these genes were also identified in susceptible strains for the same drugs, suggesting they may not be sufficient to confer the observed phenotype (**Supplemental Data 7**).

Finally, we also found that there was no significant positive correlation between AMR and the virulence phenotypes of the clinical isolates, in line with previous reports^56^.

### Genotype-phenotype correlations

To identify genotype-phenotype correlations, we performed a microbial genome-wide association study for each of the four virulence phenotypes using Scoary2, our pangenomic presence/absence gene matrix, and our phenotypic data. Cytotoxicity (4hr and 24hr) showed a statistically significant association with the presence of *exoU* (**Supplemental Data 4**), which has been documented to be the primary contributor to mammalian cell cytotoxicity^45,52,57,58^. Correlation of swimming motility, biofilm formation, and pyocyanin production to gene content did not yield any significant gene associations, likely because of the genetic complexity of these mechanisms (**Supplemental Data 3**). More generally, comparison of our phenotypic data to our phylogenetic data showed that only cytotoxicity and biofilm formation are segregated based on phylogeny (**Supplemental Figure 3A–C**).

We noted that while most strains in Group B showed high acute cytotoxicity, as evidenced by low host cell survival, 6 strains showed greater than 70% host cell survival at this time point (**Figure 6**). The lack of cytotoxicity was still evident at 24 hrs, demonstrating that this phenotype was not time-dependent (**Figure 7**). Analysis of the syntenic region encoding *exoU* demonstrated genetic rearrangement and deletion of *exoU* in BWH043; no other genes were found elsewhere in the genome with >70% homology to *exoU*, suggesting that this strain is not cytotoxic because it does not carry the *exoU* gene (**Figure 6A**). BWHPSA026 showed no evidence of acute cytotoxicity (4hr survival mean > 100%) (**Figure 7**) although its nearest phylogenetic neighbor, PS50, shows typical cytotoxicity for Group B (4hr survival mean = ∼25%). These strains encode identical *exoU* genes and differ by only 132 missense variants in protein coding genes (**Supplemental Figure 5A**). The missense mutations present in BWHPSA026 and absent in PS50 do not seem to explain the difference in cytotoxicity (**Supplemental Data 8**).

**Figure 6.**
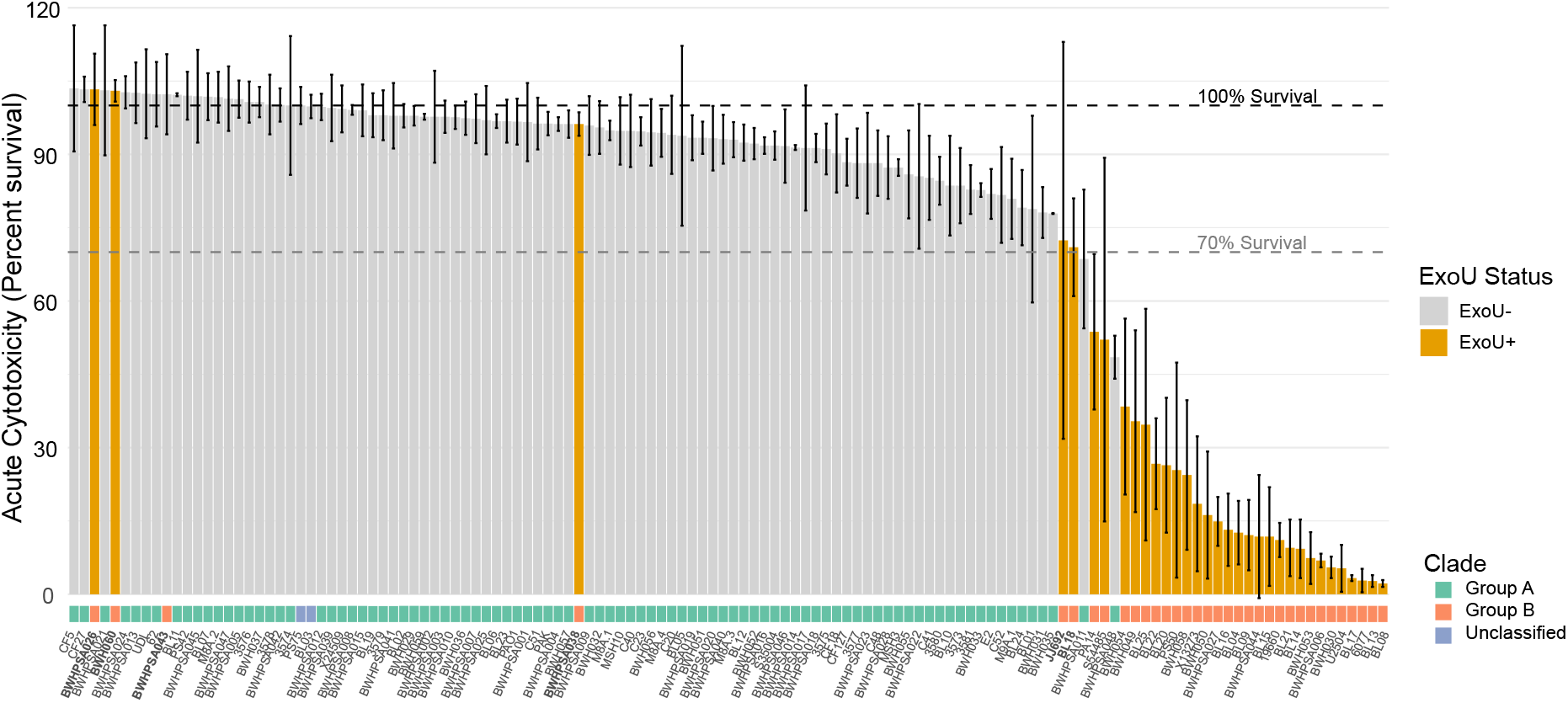
Association of cytotoxicity with *exoU* and clade. Acute cytotoxicity shown as percent survival of host cells per strain in the absence (gray) or presence (orange) of *exoU*. Clade designation shown below. Bolded strains are those that belong to Group B but show >70% mean host cell survival.

**Figure 7.**
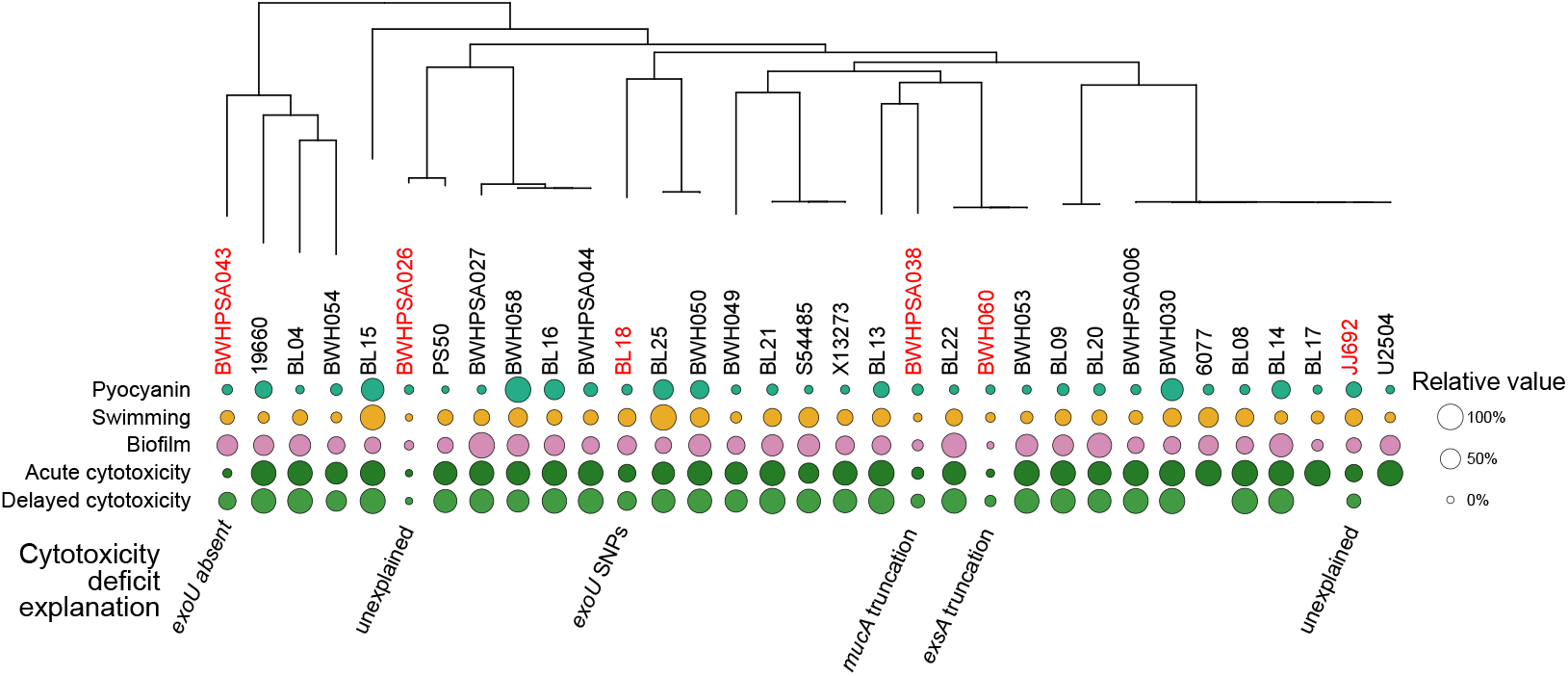
Genetic mutations of strains in phylogenetic Group B likely explain the non-cytotoxic phenotype. Six Group B strains (red) showed more than 70% acute host cell survival in the cytotoxicity assay. This attenuated cytotoxicity phenotype is still evident at 24 hours. We were able to provide potential explanations for the reduced cytotoxicity in four of these strains. For strains BWHPSA038 and BWH060, a deficit was seen across all tested virulence assays, which led to the identification of SNPs in the master regulators MucA and ExsA, respectively. Missing 24hr cytotoxicity values for strains U2504, BL17, and 6077 indicate complete host cell killing before this time point.

Sequence analysis showed that the *exoU* allele of JJ692 and BL18 had SNPs compared to the PA14 *exoU* allele (**Supplemental Data 5**). The JJ692 SNP (Pro447Leu) is also found in other strains that were cytotoxic and therefore does not explain the decreased cytotoxicity (**Supplemental Data 5**). Overexpression of the BL18 *exoU* allele in the PA14Δ*exoU* background resulted in lower cytotoxicity compared to overexpression of the WT PA14 *exoU* allele, suggesting that these point mutations may affect the ExoU phospholipase activity and, in isolation, result in lower host cell death (**Supplemental Figure 5B**). These five BL18 SNPs have not been previously described and cluster around the active site residue (Asp344) required for lipase activity^58^, suggesting that they may affect protein function (**Supplemental Figure 5C**).

The remaining 2 strains (BWHPSA038 and BWH060) encoded WT copies of *exoU*. We noted that these strains were also defective in the other acute virulence phenotypes tested (pyocyanin production, swimming motility, and biofilm production, **Figure 7**), which suggested that they may carry a mutation in a master regulator controlling all these phenotypes. In fact, we found unique mutations, not found in other strains in our collection, that could be responsible a loss of all virulence phenotypes: BWHPSA038 has a frameshift mutation (Ala144fs) in the anti-sigma factor MucA resulting in a truncated protein that lacks 51 amino acids at the C-terminus, and BWH060 has a premature stop codon (Gln43*) for the type III secretion transcriptional activator ExsA (**Supplemental Figure 5D**). The *mucA* frameshift mutation has been previously identified in mucoid strains as the *mucA22* allele, and a similar deletion (*mucA*ΔG440) was shown to result in repression of the T3SS^59,60^. Analysis of the functional domains of ExsA has demonstrated that the C-terminal domain, missing in the BWH060 strain, is necessary and sufficient for activation of type III secretion promoters^60^.

## Discussion

Comparative bacterial genomics to correlate genomic content with virulence or AMR can uncover novel mechanisms that drive pathogenic processes, especially for diverse species like *Pseudomonas aeruginosa*. Here, we performed whole genome sequencing of 125 clinical and environmental isolates, phenotyped them with four virulence assays and determined AMR phenotypes to eight antibiotics. The high variability of phenotypes amongst clinical and environmental strains demonstrates the diversity within the population. Moreover, the low level of correlation between isolation source and phenotype suggests that none of these *in vitro* phenotypes are unconditionally required for infection of a particular host tissue. Despite varied virulence phenotypes across the population, cytotoxicity was the only phenotype with a statistically and biologically meaningful gene set by a bacterial GWAS, and for the trait of acute cytotoxicity, the presence of *exoU* in particular drove sharp differences. The role of the T3SS effector ExoU in cytotoxicity has been well-documented^46,47,52,58^. Nevertheless, our analysis did uncover a novel *exoU* variant that we demonstrated yields reduced protein activity. Correlation of other phenotypes to gene content did not reveal novel pathways, likely because those phenotypes are multifactorial, controlled by many mechanisms. Our results suggest that, given the genetic diversity within the *P. aeruginosa* population, such analysis will likely require thousands, rather than hundreds, of strains.

The observation that *P. aeruginosa* has a large genome resulting from high genetic complexity, likely allows for the adaptation to diverse ecological niches^3^. We analyzed accessory genome content across isolation sources in our collection to identify genetic loci that may confer a predisposition to a given type of infection. However, we did not identify any genomic regions that uniquely correlated with the isolation source. Similarly, the virulence phenotypes we measured generally did not correlate with isolation sources. Limited exceptions were seen with environmental isolates that showed significantly lower cytotoxicity and greater motility than a subset of clinical isolate groups, and absent AMR in environmental isolates compared to clinical isolates. Notably, our results do not align with previous data suggesting that host-adaptation results in genome reduction^61^. It is important to note that the clinical data associated with the clinical strains in our collection does not differentiate between long-term versus short-term infections. Therefore, it remains possible that long-term adaptation to a mammalian host does result in genome reduction, as has been found in people with Cystic Fibrosis^62^. Altogether, our results show that disease-causing strains are not distinctly genetically dissimilar from those found in the environment.

Our analysis also allowed us to correlate virulence phenotypes and antibiotic susceptibility. Biofilm formation and motility have been previously correlated and it has been hypothesized that movement towards and attachment to a surface are required for biofilm formation^53,54^. However, we did not see a positive correlation between biofilm formation and flagella-dependent swimming motility. This is in line with a recent analysis using an independent collection of strains that also did not see a correlation between biofilm formation and any form of motility (swimming, swarming, twitching)^56^.

T3SS has a critical role in pathogenesis as it is the major contributor to mammalian cell cytotoxicity, enabling bacteria to directly inject effector proteins into the cytoplasm of host cells. The mechanisms by which ExoU, ExoT, ExoS and ExoY cause cell death have been widely studied and shown to target both the membrane and the cytoskeleton. PAO1 and PA14 are laboratory reference strains and have been used to examine the role of these effector proteins in various settings^63,64^. Both strains carry the *exoT* and *exoY* genes, while only PAO1 carries the *exoS* gene and only PA14 carries the *exoU* gene. As had been previously reported, the C-terminus of ExoY in PA14 is longer than in PAO1^49^. We found that this longer allele was only found in 4 additional strains within our collection. We also noted that the PAO1 allele of *exoT*, which encodes for a shorter protein than the PA14 allele, is much more common within our collection. This suggests that the PAO1 alleles of *exoY* and *exoT*, rather than the PA14 alleles, should be considered the WT versions and used as a reference when studying the mechanisms of action of these Exo proteins.

Given the rise in AMR, identifying the mechanisms that give rise to resistance has been a significant goal in the past decades. The ability of next-generation sequencing to identify novel genes based on, for example, sequence similarity, has revolutionized our knowledge of resistance elements and their diversity^65^. Classification of AMR by sequence analysis remains imperfect, and we must still rely on laboratory-based susceptibility testing, which is both costly and time-consuming since it must be done for each antibiotic individually. Prior genomic analysis of 390 *P. aeruginosa* strains demonstrated that susceptibility to meropenem and levofloxacin could be readily determined from sequence analysis, but determining susceptibility to amikacin was less tractable^66^. In our study, we were able to assign genetic determinants of resistance in ∼93% of cases where intermediate resistance or resistance was identified, demonstrating a high probability of sequenced-based susceptibility classification for the antibiotics that we analyzed. As more strains are sequenced and more mechanisms of resistance are identified, sequenced-based susceptibility classification should become more feasible and reliable. Lastly, a more detailed understanding of mechanisms of resistance may point to novel ways to subvert these pathways, potentially allowing us to continue to rely on current antibiotics.

In conclusion, our comprehensive genomic and phenotypic analysis of 125 clinical and environmental *P. aeruginosa* isolates reveals that this opportunistic pathogen’s remarkable versatility stems from a broadly conserved genomic repertoire rather than niche-specific gene acquisition. The lack of strong genotype-phenotype correlations for most virulence traits, along with the absence of isolation source-specific genomic signatures, highlights *P. aeruginosa*’s capacity for opportunistic colonization across diverse environments and host tissues. The apparent dichotomy between AMR patterns in clinical vs. environmental strains underscores the critical role of antibiotic selection pressure in driving resistance evolution. The identification of novel *exoU* variants with altered cytotoxicity demonstrates that functional diversity within well-characterized virulence factors may still influence pathogenic outcomes. These findings taken together bolster our understanding of *P. aeruginosa* pathogenesis and evolution, and our rich genotype-phenotype dataset generated here provides a valuable resource for future studies investigating the pathogenic potential of this versatile species.

## Acknowledgments

We thank members of the Hung lab for helpful discussion and Roberto Kolter, Paula Suarez, Wolfgang Haas (Bausch & Lomb), Fred Ausubel, and Steve Lory for sharing their strain collections. Strains with BWH or BWHPSA nomenclature are from the Crimson Biomaterials Collection Core Facility at Mass General Brigham, Boston, MA. This work was funded by grants to DTH (1R21AI097613-01) and JR (1U01AI176414), and institutional funds to CP (National Jewish Health) and JR (start-up funds and Translational Research Scholars Program from the University of Colorado Anschutz Medical Campus).

## Supplementary data

**Supplemental Data 1. Genome accession IDs and associated metadata**.

Sheet *SRA_IDs* contains the strain accession numbers and sequencing metadata for our 125 isolates. *Ref_Strain_IDs* lists the RefSeq accession numbers for laboratory reference strains used. *Strain_origins* shows the original isolation source metadata recorded (*PreciseSource*) as well as our fine classification (*Source*) and broad classification (*GroupedSource*). *Phenotypes* records the antimicrobial susceptibility testing results for tested compounds, plus the mean, standard deviation, and replicates for the virulence assays performed.

**Supplemental Data 2. Tree metrics**.

Sheet *ModelFinder_Results* shows the best performing substitution models as chosen by ModelFinder for our dataset. AIC and BIC yielded slightly different best models, and for our set we selected the top model based on BIC. *IQ-TREE2_ParameterResults* reports the initial and final model parameters for our phylogenetic estimation.

**Supplemental Data 3. Consensus phylogenetic tree with confidence estimates**

We provide the unmodified output consensus tree built using core SNPs with estimated confidence values from IQ-TREE2 as described above.

**Supplemental Data 4. Pangenomic associations with trait and isolation source**.

A microbial genome-wide association study was performed with Scoary2 to associate genes with virulence traits or isolation sources. As an example, *4hr_Cytotox_Scoary2* reports *exoU* as a top hit for the trait of host cell survival – that is, g-(lacking genotype *exoU*) is associated with t+ (trait of greater host cell survival). Few associations across all tests remained significant after controlling for multiple testing. Many genes have uninformative “group_####” names, indicating that consensus names were not identified during pangenome feature clustering. Representative protein sequences are provided for downstream investigation of all features.

**Supplemental Data 5. Sequences and polymorphisms for exoenzymes**.

*Summary_nonSYN* lists non-synonymous mutations in *exoU, exoT, exoS*, and *exoY*; WT indicates same sequence as reference, “-” indicates gene is not found in genome. *exo_AllSNPs* lists a comprehensive set of all SNPs observed in all strains, while subsequent sheets are subsetted per-exoenzyme.

**Supplemental Data 6. Correlation analysis**.

We tested whether virulence traits were correlated among isolates from different sources or phylogenetic groups. For example, *GroupA_Subset* reports Spearman and linear regression modeling tests for correlation between different traits in Group A isolates. As expected, 4hr and 24hr host cell survival are significantly correlated, while most other traits are not strongly correlated.

**Supplemental Data 7. Computationally identified AMR mechanisms per strain**.

We used AMRFinderPlus to identify mechanisms for antimicrobial resistance in our strains. We listed features found in >50% of strains “Common_mechanisms,” and rarer mechanisms “Specific_mechanisms.” If explanatory mechanisms for resistance towards a given drug were identified, they are recorded as TRUE, and if specific mechanisms are lacking, FALSE is shown. If an isolate showed susceptibility for a given drug, the mechanistic explanation for that phenotype is simply reported as NA.

**Supplemental Data 8. Differentiating SNPs in strains PS50 and BWHPSA026**.

PS50 and BWHPSA026 showed stark cytotoxicity differences despite few genetic differences (132 missense variants in protein coding genes). PS50 was cytotoxic (25% host cell survival), while BWHPSA026 was not (100% host cell survival).

