## Supplemental Figures for "Mapping genetic and phenotypic diversity of *Pseudomonas aeruginosa* across clinical and environmental isolation sites"

**Supplemental Figure 1. Genome assembly and annotation statistics.**

All whole genome sequencing data were cleaned and checked for quality using FASTP, assembled *de novo* using Shovill, and annotated using Bakta. Summary statistics overall and per isolation source are provided.

**Supplemental Figure 2. Clade classification of *Pseudomonas aeruginosa* strains.**

**A**. A SNP-based phylogenetic tree rooted on the phylogenetic outlier PA7 in comparison with the strains used in this study. **B**. Clade distribution for each isolation source. **C.** Pairwise Fisher's Exact Test results for clade distribution. Environmental strains were overrepresented in Group A, while urine and eye strains were overrepresented in Group B.

**Supplemental Figure 3. Correlation of clade and virulence phenotypes.**

Virulence phenotypes were compared between Group A (n=92) and Group B (n=34) strains. Biofilm formation and cytotoxicity (decreased host cell survival) were significantly greater in Group B strains. Wilcoxon test with Benjamini-Hochberg correction for multiple testing: $^{**}p<0.01$; $^{***}p<0.001$.

**Supplemental Figure 4. Antibiotic Susceptibility Testing Phenotypes.**

Antibiotic Susceptibility Testing phenotype for each strain-antibiotic combination based on the Zone Diameter Interpretive Chart for *P. aeruginosa* reference strain ATCC 27853.

**Supplemental Figure 5. Non-cytotoxic Group B strains.**

**A.** A heatmap showing the pairwise SNP distance between Group B strains. **B.** The BL18 *exoU* allele shows reduced cytotoxicity compared to the PA14 reference. Deletion of *exoU* from PA14 yields high host cell survival as expected, and overexpression from a plasmid vector with the PA14 *exoU* allele restores wild-type toxicity. Expression of the BL18 *exoU* allele significantly increased host cell survival, supporting the *exoU* locus as responsible for the BL18 strain's attenuated host cell killing. **C.** SNPs in the BL18 *exoU* allele compared to PA14 yield a cluster of missense mutations in the C-terminal portion of the ExoU domain (Conserved Domain Database ID: cd07207). **D.** SNPs in BWH060 *exsA* allele compared to PA14 show a truncation with a premature stop at Q43. **E.** A missense SNP in the BWH038 *mucA* allele compared to PA14 results in a frameshift at A144.

Supplemental Figure 1

| Genome assembly and annotation statistics | | | | | |
|---|---|---|---|---|---|
| | Number of Isolates | Mean Length (±SD) | Length Distribution | Mean CDSs (±SD) | CDSs Distribution |
| Overall summary statistics | | | | | |
| Total | 128 | 6.65 Mbp ± 0.28 Mbp | 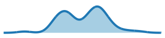 | 6074 ± 269 | 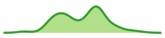 |
| Summary statistics by isolation source | | | | | |
| blood | 9 | 6.79 Mbp ± 0.15 Mbp | 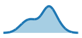 | 6212 ± 138 | 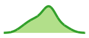 |
| environmental | 28 | 6.62 Mbp ± 0.28 Mbp | 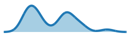 | 6050 ± 287 | 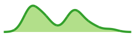 |
| eye | 27 | 6.75 Mbp ± 0.27 Mbp | 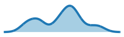 | 6150 ± 266 | 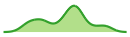 |
| reference | 3 | 6.40 Mbp ± 0.14 Mbp | 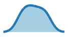 | 5830 ± 136 | 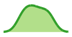 |
| other | 11 | 6.63 Mbp ± 0.23 Mbp | 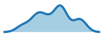 | 6080 ± 244 | 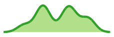 |
| pulmonary | 20 | 6.59 Mbp ± 0.30 Mbp | 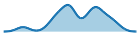 | 6014 ± 277 | 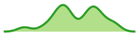 |
| urine | 22 | 6.64 Mbp ± 0.32 Mbp | 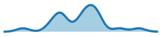 | 6048 ± 300 | 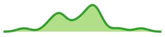 |
| wound | 8 | 6.60 Mbp ± 0.23 Mbp | 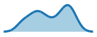 | 6043 ± 230 | 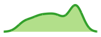 |

Supplemental Figure 2

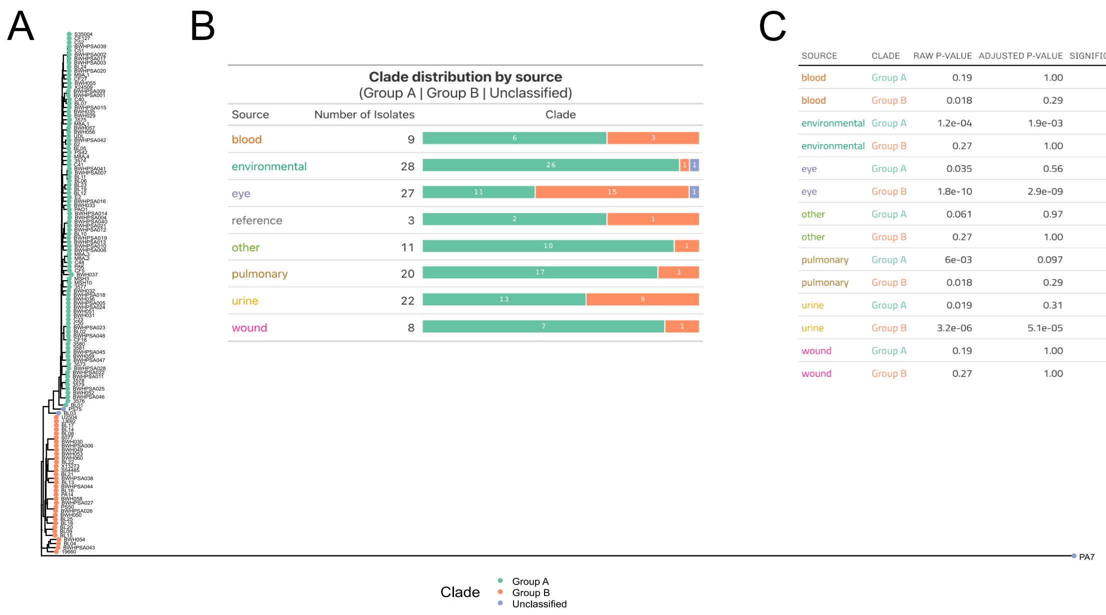


C

| SOURCE | CLADE | RAW P-VALUE | ADJUSTED P-VALUE | SIGNIFICANCE |
|---|---|---|---|---|
| blood | Group A | 0.19 | 1.00 | |
| blood | Group B | 0.018 | 0.29 | |
| environmental | Group A | 1.2e-04 | 1.9e-03 | ** |
| environmental | Group B | 0.27 | 1.00 | |
| eye | Group A | 0.035 | 0.56 | |
| eye | Group B | 1.8e-10 | 2.9e-09 | *** |
| other | Group A | 0.061 | 0.97 | |
| other | Group B | 0.27 | 1.00 | |
| pulmonary | Group A | 6e-03 | 0.097 | |
| pulmonary | Group B | 0.018 | 0.29 | |
| urine | Group A | 0.019 | 0.31 | |
| urine | Group B | 3.2e-06 | 5.1e-05 | *** |
| wound | Group A | 0.19 | 1.00 | |
| wound | Group B | 0.27 | 1.00 | |

Supplemental Figure 3

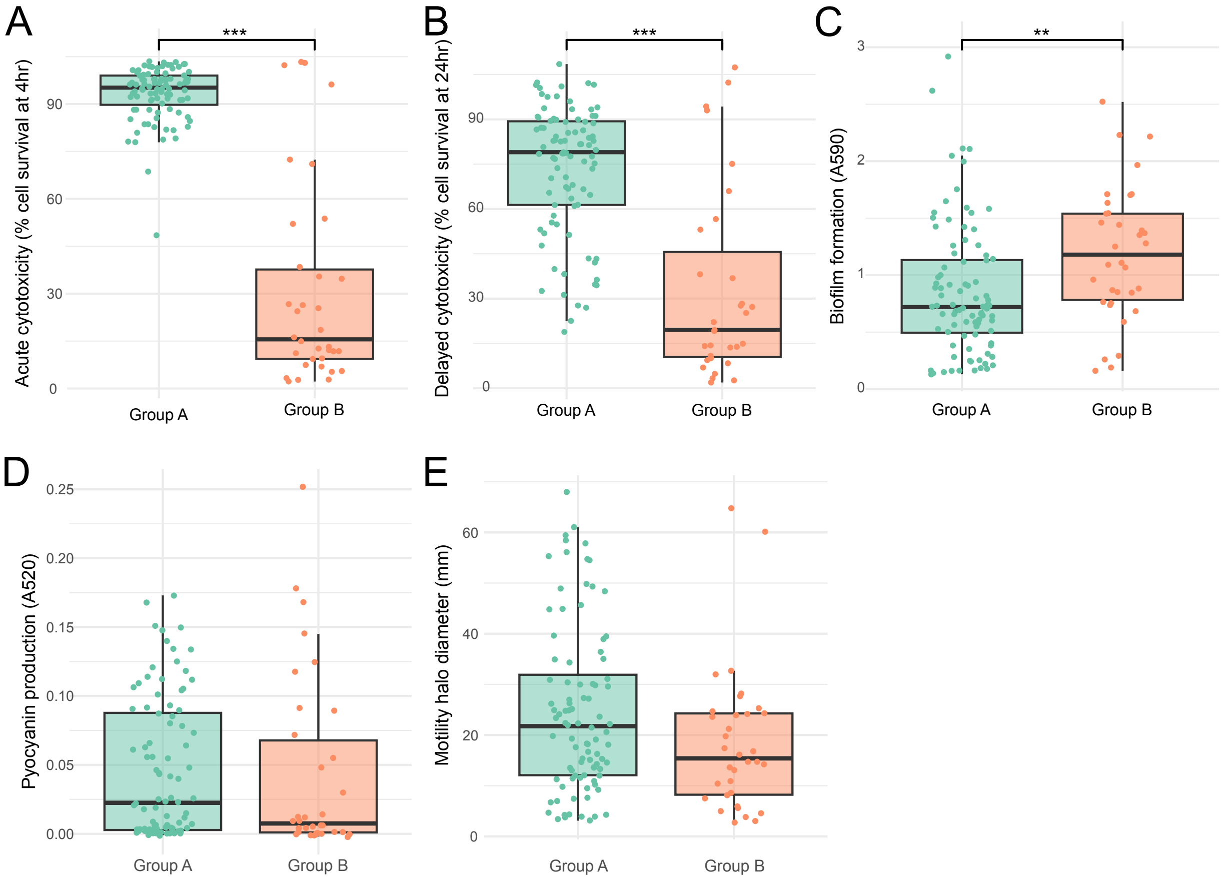

### Supplemental Figure 4

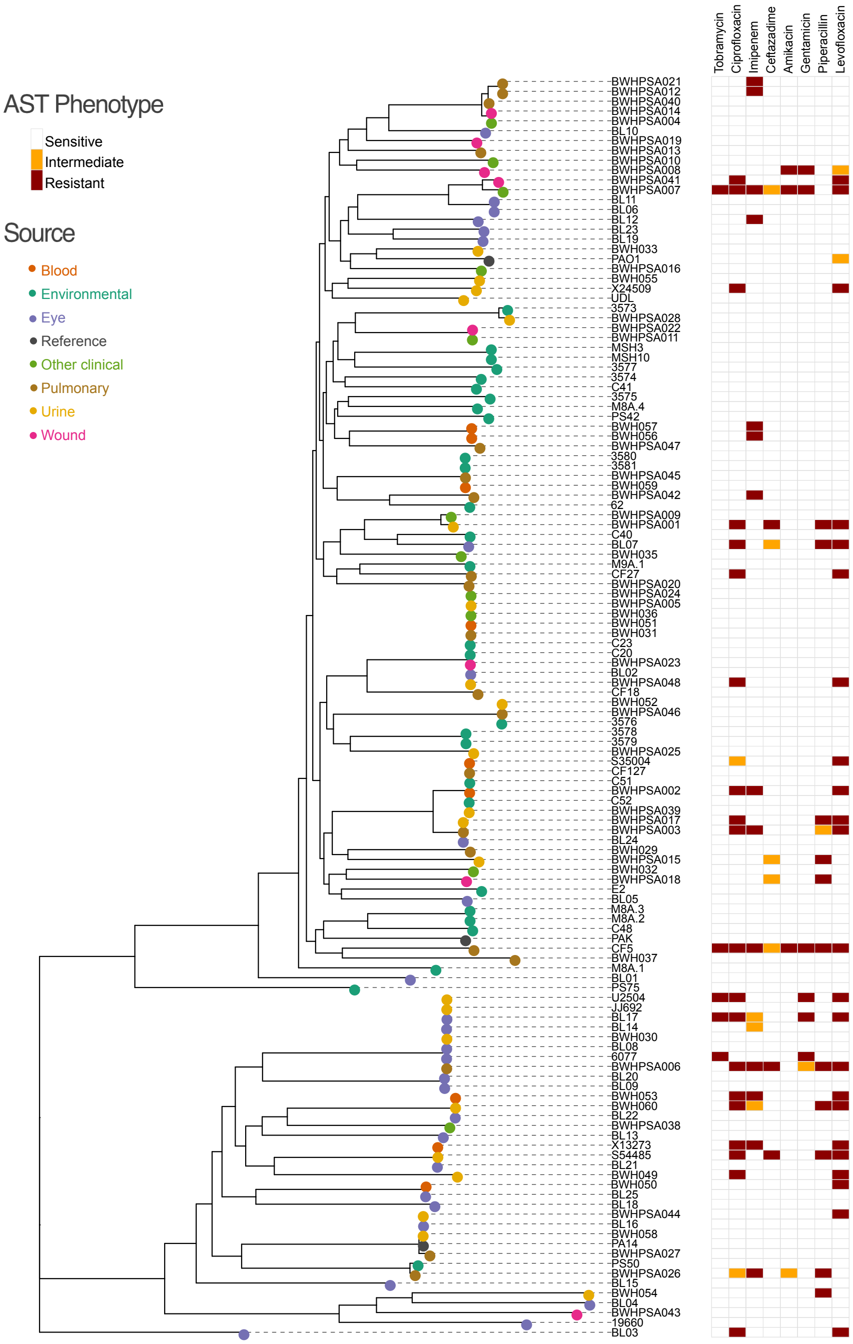

Supplemental Figure 5

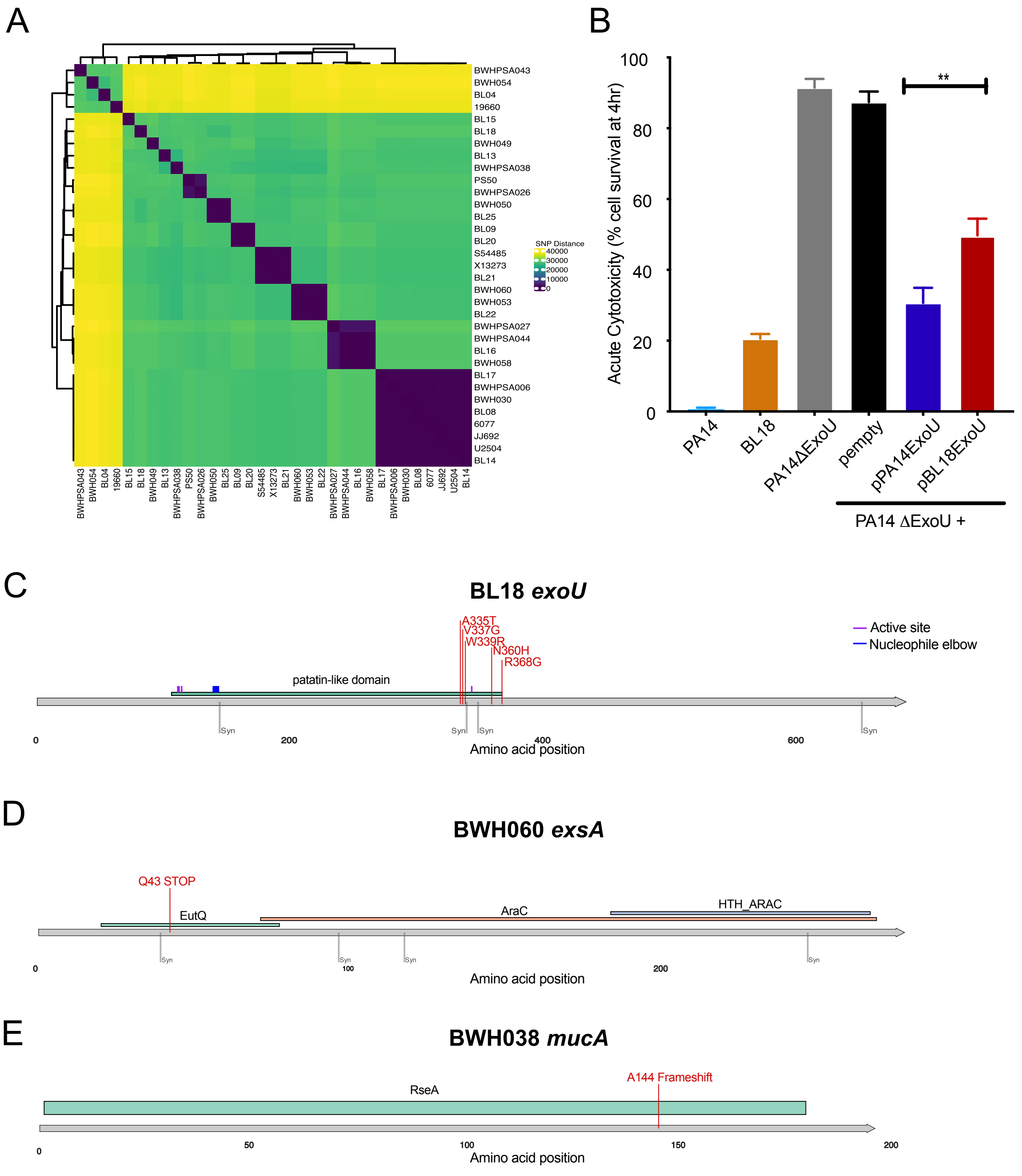
